# Role of intraflagellar transport in transcriptional control during flagellar regeneration in Chlamydomonas

**DOI:** 10.1101/2022.09.29.510156

**Authors:** Karina Perlaza, Ivan Zamora, Wallace F. Marshall

## Abstract

Biosynthesis of organelle precursors is a central part of the organelle size control problem, but what systems are required to control precursor production? Genes encoding flagellar proteins are upregulated during flagellar regeneration in *Chlamydomonas*, and this upregulation is critical for flagella to reach their final length, but it not known how the cell triggers these genes during regeneration. Here we present two models based on transcriptional repressor that is either produced in the flagellum, or else is produced in the cell body and sequestered in the growing flagellum. We show that both models lead to stable flagellar length control, can reproduce the observed dynamics of gene expression, and are consistent with the effects of protein synthesis inhibitors on gene expression. The two models make opposite predictions regarding the effect of mutations that block intraflagellar transport (IFT). Using quantitative measurements of gene expression, we show that gene expression during flagellar regeneration is greatly reduced in mutations of FLA3, FLA8, and FLA10, which encode the three components of the heterotrimer kinesin-2 that drives IFT. This result is consistent with the predictions of the model in which a repressor is sequestered in the flagellum by IFT. In contrast to the effects of IFT mutants, we find that inhibiting axonemal assembly has much less effect on gene expression, suggesting that transport is more important than axonemal assembly. The repressor sequestration model allows precursor production to occur when flagella are growing rapidly, representing a form of derivative control.

## Introduction

A fundamental question in cell biology is how cells control the size of organelles and other cellular structures. Potential models for organelle size control have focused on either the regulation of the assembly process, or the regulation of precursor synthesis (Goehring 2012; Marshall 2016; Mohapatra 2017), since size could in theory be determined by either process. In fact, both are likely to be important.

The flagella of the unicellular green alga *Chlamydomonas* provide a highly tractable model system in which to study organelle size regulation (Randall 1969; Wemmer 2007). Flagellar assembly and length maintenance involves a kinesin-based transport mechanism known as intraflagellar transport or IFT (Kozminski 1993; Cole 1998; Rosenbaum 2002; Bhogaraju 2014) that actively transports tubulin and other building blocks to the site of assembly at the tip of the growing flagellum (Qin 2004; Hao 2011; Bhogaraju 2013; Craft 2015). The activity of the IFT pathway is a function of flagellar length, such that the rate of IFT decreases according to 1/L (Engel 2009; Ludington 2013), but the mechanism by which length regulates IFT remains unclear (Ludington 2015; Ishikawa 2017; Hendel 2018; Ishikawa 2022). The transport of tubulin and other cargos by IFT also varies as a function of length (Wren 2013; Craft 2015), but at least in the case of tubulin it remains unclear whether this represents regulation of a binding interaction or a length dependence of the number of binding sites (Wemmer 2020). Because flagellar microtubules undergo constant disassembly at steady state (Marshall 2001; Song and Dentler, 2001), it is thought that the balance of IFT-mediated assembly and constant disassembly sets the steady state length (Wemmer 2007). Because IFT is a decreasing function of length (Engel 2009; Ludington 2013), when flagella regenerate, they initially grow rapidly, but then the growth rate decreases as the flagella approach their final steady-state length (Marshall 2005) thus producing a stable steady state solution for length.

Flagellar assembly depends not just on IFT, but also on the availability of precursor proteins to transport in the first place. When *Chlamydomonas* flagella are removed, for example by pH shock, they grow back in approximately an hour to the length they had before removal. If, however, protein synthesis is inhibited, flagella only grow back to half their normal length (Rosenbaum 1969). This result indicates that cells contain enough protein to build half-length flagella, but that further growth requires new protein synthesis.

Consistent with this observation, it has been found that genes encoding flagellar proteins are upregulated during flagellar regrowth (Schloss 1984; Lefebvre 1980; Baker 1984; Lefebvre 1986). This fact has been exploited to identify flagella-related genes based on their upregulation during flagellar assembly (Stolc 2005; Pazour 2005; Chamberlain 2008; Albee 2013; Lin 2015; Zones 2015). These studies have shown that for most flagellar genes, mRNA levels increase over a time-scale of approximately 30 minutes, and then decrease back to baseline over the next hour, creating a pulse of gene expression that coincides with the growth of the flagella. The increase is due to alteration in transcription, as evidenced by the fact that mutation of the promotor of upregulated genes can prevent their upregulation (Davies 1994).

The pulse-like wave of flagellar gene expression during flagellar regeneration raises the question of what is the signal that regulates this pulse? How might flagellar loss or regeneration serve as a trigger of gene induction? It has been proposed that a positive regulator of transcription might be produced or activated when flagella regenerate. For example, one proposal is that calcium influx that occurs when flagella detach might trigger gene expression (Evans 1997). An alternative possibility is suggested by examples from the context of animal regeneration, in which it has instead been found that negative regulators often are the basis for controlling regeneration pathways. One classic example is the regeneration of the eyestalk in crustaceans, in which a hormone produced within the eyestalk inhibits the regeneration pathway by acting on a second region in the brain (Mykles 2021). When the eyestalk is severed, the source of the inhibitory hormone is removed, and this leads to induction of regeneration. In contrast to the eyestalk example, flagellar detachment is not required to change gene expression in *Chlamydomonas*. When full-length flagella are induced to elongate or shorten, expression of flagellar genes increases or decreases, respectively (Periz 2007; Chamberlain 2008). These data suggest that some factor related to flagellar length or growth may play a role in generating the necessary signal.

Here we consider the possibility that flagella-related gene expression is under control of a negative regulator, either in the form of an inhibitory signal produced by the flagellum, or a repressor molecule that is synthesized in the cell body and sequestered in the growing flagellum. We will refer to these two models as the repressor production and repressor sequestration models, respectively. The latter model was first proposed by Lefebvre and Rosenbaum (1978), who envisioned a repressor of protein synthesis that assembles into the growing flagellum. Although the Lefebvre and Rosenbaum model was focused on translational control, the same idea would apply at the level of transcriptional regulation. We show that both the repressor production and repressor sequestration models can produce stable flagellar length control accompanied by pulsatile gene expression during flagellar regeneration. Using conditional mutants in the heterotrimeric kinesin-2 that drives IFT, we find that impaired flagellar regrowth and IFT leads to lower peak expression of flagella-related genes in a manner consistent with predictions of the repressor sequestration model but not the repressor-production model.

## Results

### Dynamics of flagellar gene induction relative to flagellar growth

Flagellar regeneration can be induced by a number of stresses, but pH shock (Schloss 1984) is particularly convenient (**Figure 1A**). In this procedure, cells grown in neutral pH media are exposed to a reduced pH for one minute by addition of acetic acid, and then restored back to pH 7 by the addition of KOH. During the shock, flagella detach via an active severing process (Quarmby 2004) and then regenerate over approximately an hour (Rosenbaum 1969). During this time, nuclear genes encoding flagellar proteins are upregulated. Eventually, as flagella reach their final steady state length, mRNA levels drop back down to the pre-shock level. By measuring flagellar length after pH shock together with mRNA levels for the flagellar protein RSP3 measured by quantitative PCR (qPCR) as described in Materials and Methods, we see that there is a clear pulse of gene expression whose peak coincides with the time at which the flagellum is growing the most rapidly. The mRNA level is proportional to the slope of the flagellar growth curve (data not shown). Expression levels begin to drop as the flagellar growth rate decelerates. In **Figure 1C**, the same data are plotted except that the flagellar growth rate (dL/dt) is plotted alongside the mRNA levels. Comparing **Figures 1B and 1C**, it can be seen that mRNA levels are not correlated with flagellar length, but with the flagellar growth rate.

**Figure 1.**
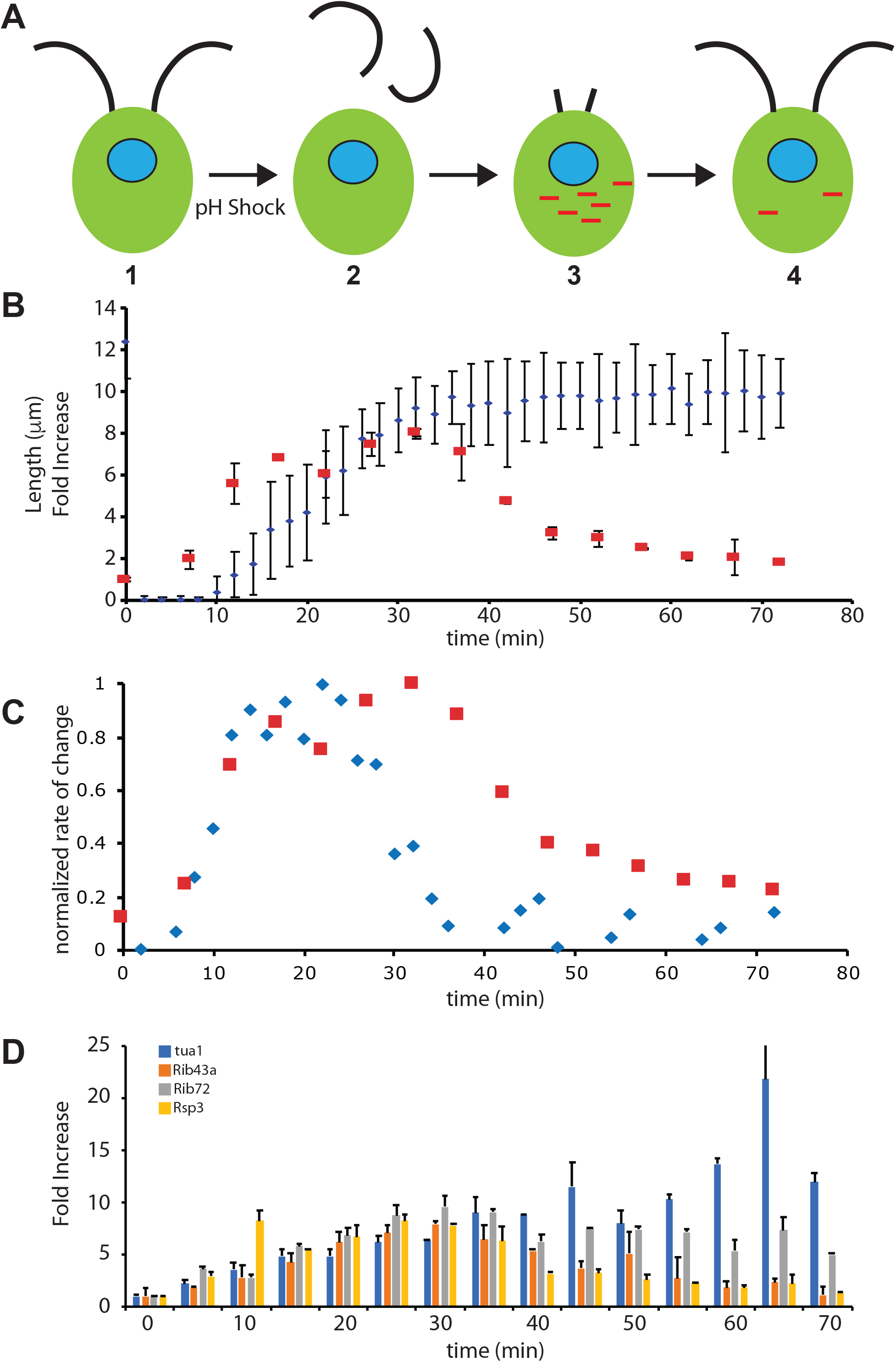
Transcriptional dynamics of flagella-related genes relative to flagellar growth. (**A**) Flagellar regeneration following pH shock. (1) During normal growth, *Chlamydomonas* cells have two full-length flagella, and transcription of flagella-related genes is at basal levels. (2) Transient pH shock causes flagella to detach. (3) As flagella begin to regrow, genes encoding flagellar proteins in the nucleus (blue) are transcribed, leading to accumulation of mRNA (red). (4) As flagella reach final length, transcription returns to basal levels. (**B**) Flagellar length (blue) and fold induction of RSP3, a flagella-related gene (red), based on qPCR, as a function of time after pH shock. Error bars are standard error of the mean for flagellar length, and standard deviation between biological replicates for qPCR. (**C**) fold induction of RSP3 (red) plotted along with flagellar growth rate (blue) based on the data of panel B. Both quantities were normalized to their maximum values. (**D**) Timecourse of expression of four flagella-related genes during regeneration based on qPCR. All four genes show a parallel increase up to the peak of expression at 30 minutes.

The correlation of mRNA levels with flagellar growth rate is not restricted to the RSP3 gene. As shown in **Figure 1D**, we see very similar induction profiles for genes encoding alpha tubulin, the axonemal structural proteins Rib43a and Rib72, and radial spoke protein 3 (Rsp3) which is part of the motility machinery. Most of these genes followed roughly similar kinetics in their rate of increase, just as has been previously reported for other genes (Schloss 1984; Baker 1984; Remillard 1982; Chamberlain 2008), but vary in the duration of the pulse, with tubulin showing a more sustained elevation of message level. These data suggest that induction of the genes may be taking place with similar kinetics, but then the different mRNA species may vary in their stability once expression ceases. The time course of expression that we see here is consistent with a previous qPCR study that included several of the same genes (Chamberlain 2008).

Given that these diverse genes appear to be turned on with similar kinetics, we ask what might be the stimulus that triggers their expression.

Given that the process of flagellar regeneration was initiated by a pH shock, one trivial possibility might be that the stress of the pH shock itself is the trigger. In this case, flagellar growth would be irrelevant and the correlation of mRNA with growth rate would just be a coincidence. To test this possibility, we measured gene induction by pH shock in the *bld1* mutant of Chlamydomonas, a mutation in IFT52 that lacks flagella (Brazelton 2001). Compared to wild-type cells (**Figure 2A,B**), bld1 mutants show a negligible change in gene expression (**Figure 2C,D**). A similar result was obtained in *bld1* mutants using Nanostring detection to assay mRNA levels for a panel of 17 induced genes including tubulin, IFT proteins, and axonemal proteins (**Figure 2E**). A similar lack of induction was seen in the *bld10* mutation (**supplemental figure S1A**), which lacks flagella due to an absence of basal bodies (Matsuura 2004).

**Figure 2.**
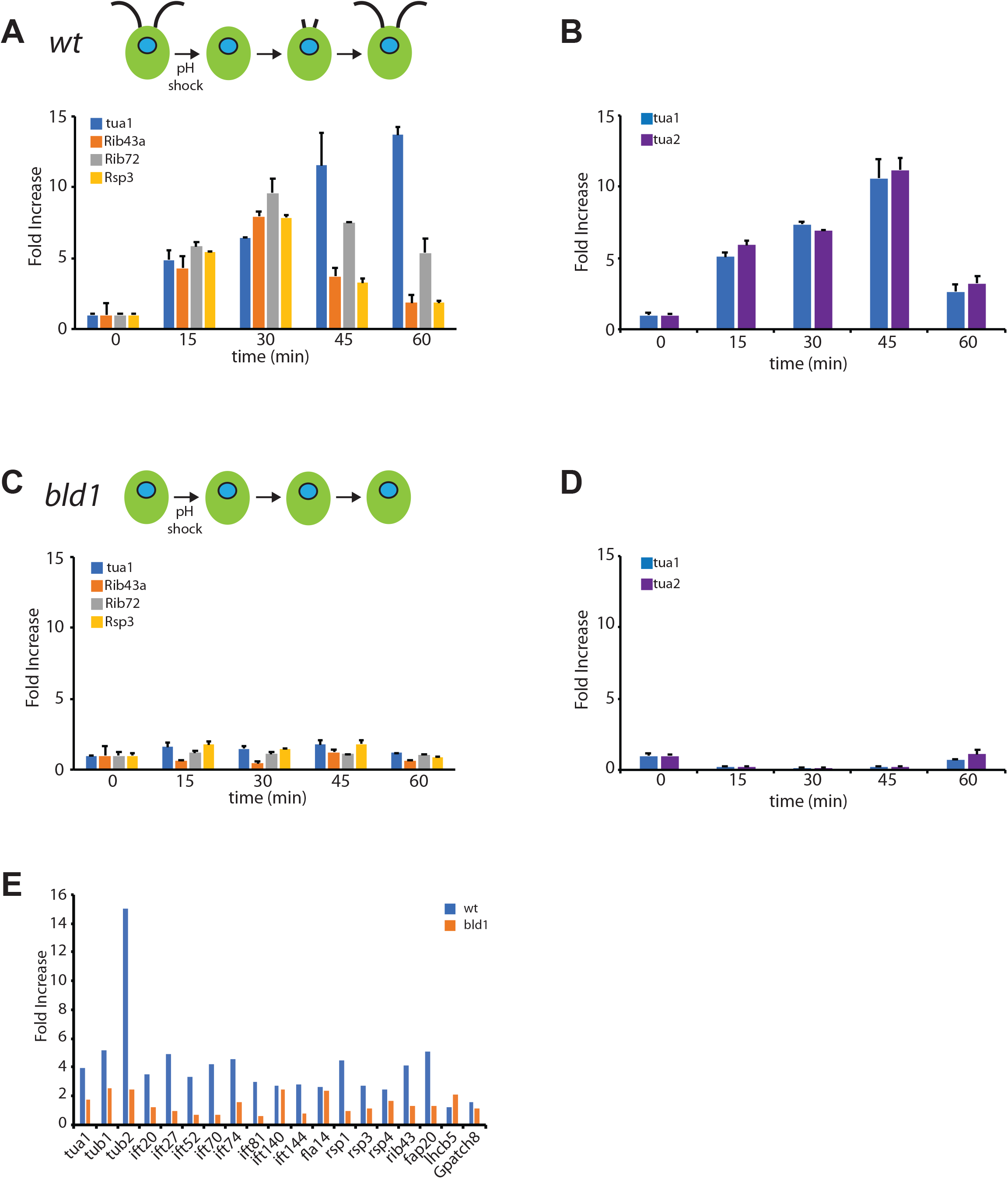
Flagellar gene expression is not triggered by pH shock. (**A**) Expression of tubulin (tua1), Rib43, Rib72, and Rsp3, in wild type cells, at four time points during regeneration analyzed by qPCR, taken from Figure 1C. (**B**) Expression of two tubulin genes during regeneration at the same time points as in panel A, from a separate experiment analyzed by qPCR. (**C**) Expression of flagella-related genes following pH shock in the *bld1* mutant, which lacks flagella. Compared to panel A, expression is drastically reduced and is comparable to basal levels at the t=0 time point. (**D**) Expression of tubulin genes following pH shock in *bld1* mutants, again showing failure to induce expression. (**E**) Expression of a panel of flagella related genes measured 30 minutes after pH shock by Nanostring detection. (blue) wild type cells, (orange) *bld1* mutants. Lhcb5 and gpatch8 are non-flagellar genes included as controls.

These results show that flagellar gene induction is not a direct consequence of the pH shock itself, but instead must be a response either to loss of flagella or to flagellar regeneration.

### A repressor-sequestration model for flagellar gene regulation

We consider two possible models for regulation of gene induction as a function of flagellar assembly. In the first model (**Figure 3A**), we envision that some component of the assembled flagellum produces a negative regulator that traffics to the cell body and inhibits gene expression, such that as long as a full-length flagellum is present, flagellar genes are repressed. In this model, the loss of flagella is sensed by the loss of the repressive signal. In the second model (**Figure 3B**), we posit a repressor (or an upstream signaling molecule that eventually stimulates a repressor) that is sequestered inside the growing flagellum, leading to induction of genes during the period of rapid flagellar growth, and such that as the flagellum reaches its final length and growth slows down, the repressor is able to accumulate in the cell body and repress those same genes. In this model, the cell does not detect loss of the old flagellum, but rather, the regrowth of the new flagella. We will consider both of these models and ask if they can, at least in principle, account for the pulse of gene expression that accompanies flagellar regeneration, while also leading to maintenance of a stable flagellar length.

**Figure 3.**
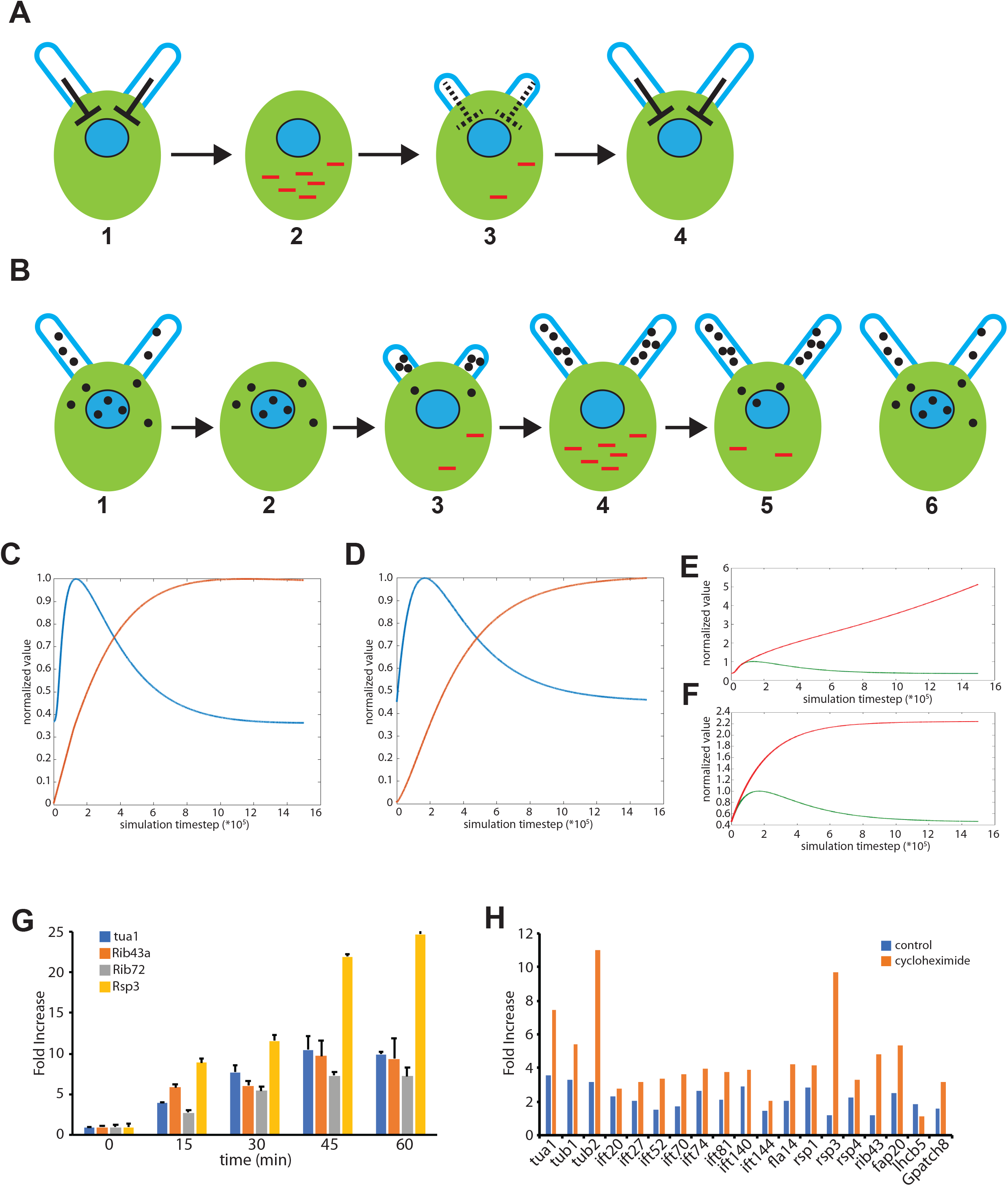
Models for regulation of flagellar genes. (**A**) Repressor Production Model in which a repressive signal, indicated by the bar-end arrows, is produced within the flagellum proportional to the current flagellar length. Panel 1 depicts the situation in cells with full length flagella. In this case, the flagella produce a repressor which keeps gene expression of flagellar genes turned off. (2) when flagella are removed, the repressive signal produced by the flagella is eliminated, leading to an increase in gene expression. (3) as flagella begin to grow back, the repressive signal starts to be produced, leading to gradual reduction in transcription. (4) when flagella reach full length, the repressive signal is maximal, leading to minimal gene expression. Overall, this sequence of events should lead to a pulse in gene expression that coincides with the early stages of flagellar regrowth. (B) Repressor Sequestration Model in which a repressor is synthesized in the cell body and transported into growing flagella by IFT. (1) in cells with full length flagella, the repressor is found in both the flagellum and in the cell body. The repressor molecules in the cell body keep gene expression at a basal level. (2) when flagella are removed, there is no immediate gene expression because repressor is still present in the cell body. (3) as flagella start to regrow using the existing pool of precursor protein, repressor starts to be imported into the growing flagella, leading to a drop in repressor concentration in the cell body. This leads to an increase in gene expression. (4) As flagella grow, more and more repressor is imported into the flagella, leading to a further increase in gene expression. Among the genes expressed is that encoding the repressor itself. (5) As flagellar growth decelerates, import of repressor into the flagella slows down due to the 1/L dependence of IFT mediate trafficking. This slowdown in import, combined with the expression of the repressor gene itself, leads to an increase in repressor concentration inside the cell body, which begins to reduce gene expression. (6) By the time flagella reach full length, repressor has accumulated back to its pre-shock concentration inside the cell body, causing gene expression to drop back to basal levels. As with the repressor production model, this repressor-sequestration model should be able to produce a pulse of gene expression. (**C**) Numerical simulation of flagellar regeneration based on repressor production model, equations A1–A4. (blue lines) predicted mRNA levels as a function of time. (red lines) predicted flagellar length as a function of time. Both flagellar length and mRNA levels are plotted normalized to their maximum value in each simulation. (**D**) Numerical simulation of flagellar regeneration based on repressor sequestration model, equations B4–B6. (**E**, **F**) Simulations of flagellar regeneration in the absence of protein synthesis based on the repressor production (E) or sequestration (F) models. In the repressor production model, mRNA levels continue to increase because the flagellum does not reach full length and thus cannot establish full production of repressor. In the repressor sequestration model, mRNA levels continue to increase because repressor protein is not synthesized. (**G**) Expression of flagella related genes in cycloheximide treated cells as a function of time after pH shock, measured by qPCR, showing increased expression levels at later timepoints. (**H**) Quantification of flagella-related genes 140 min after pH shock by Nanostring detection, showing sustained increased gene expression in the presence of cycloheximide. The non-flagellar control genes do not show this effect.

### Model of repressor produced by flagellum

We begin with the repressor production model, in which the flagellum produces a repressor at a rate proportional to its length (**Figure 3A**). This mechanism would be in effect if some signal-generating molecule, such as a kinase, was distributed along the axoneme, and then phosphorylated the actual repressor, putting it into an active state. In order to test whether this model could be plausible, we developed a coarse-grained model for flagellar gene control coupled with flagellar growth and turnover. We describe the state of the cell using three state variables, M, P, and L, which correspond to the quantity of mRNA and protein (M and P) of a flagellar regeneration-induced gene, and the length L of the flagella. In this simplified version of the model we assume that all flagellar genes are regulated with identical dynamics. We therefore model just a single protein and assume that the quantity of all structural proteins will be the same. We denote this quantity P and express it in length-equivalent units, that is, P=1 corresponds to the quantity of precursor used to build a flagellum of length 1.

We model mRNA synthesis as occurring at a constant basal rate which can be reduced by binding of repressor via first order saturable binding to the regulatory region of each flagellar gene.

Protein synthesis is assumed to occur at a rate proportional to message level. Protein degradation is assumed to occur with first order kinetics, but it is further assumed that only free protein in the cytoplasm is subject to degradation, while protein incorporated into the flagellum is not.

Changes in flagellar length follow the previously described “balance-point model” in which flagella grow at a rate proportional to the available protein precursor pool and to 1/L (Ludington 2015) and disassemble at a constant length-independent rate D. This model assumes that cargo protein is recruited from the cytoplasm and associated with IFT particles via a first order binding process operating in the linear range, so that the level of cargo loading onto a given IFT particle is directly proportional to the free protein pool in the cytoplasm. We assume a cell has two flagella, to match the situation in *Chlamydomonas*, and we express protein levels in length-equivalent units.

We denote the total pool of flagellar precursor protein as P, which includes both the protein already incorporated into the two flagella as well as the remaining cytoplasmic pool of protein.

Differential equations describing change in mRNA level (M), flagellar length (L) and cytoplasmic precursor protein level (Y) are given as follows:

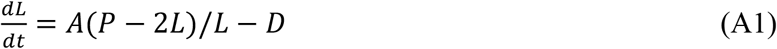

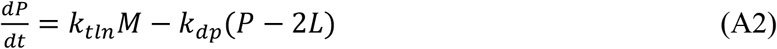

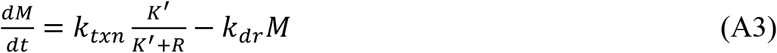

K’ describes the dissociation constant for binding of the repressor not only to the promotor of genes encoding flagellar proteins

We model the dynamics of the repressor, R, by assuming first order decay and production at a rate proportional to the instantaneous flagellar length:

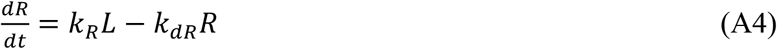

As derived in Materials and Methods, this system of equations has a unique steady state solution with a positive length equal to:

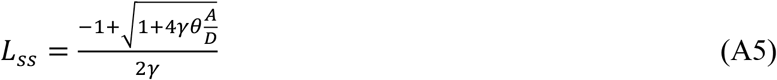

Where we have defined lumped parameters

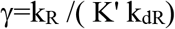

Which represents the efficacy of the repressor

and

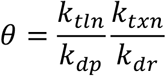

Which represents the combined processes of transcription and translation Thus a non-zero steady state solution for flagellar length always exists given the way our parameters are defined. Consistent with intuition, increasing the constant A describing IFT activity leads to increased flagellar length, as does decreasing the flagellar disassembly rate D. Increasing the affinity of the repressor to the promotors (decreasing K’ and thus increasing g) predicts a decreased length, again as expected since precursor production would be shut off at shorter lengths.

We can use these equations to model the predicted dynamics of gene induction during flagellar regeneration, by setting the length near zero to mimic flagellar detachment, and reducing the protein pool by the equivalent of two full length flagella, and then solving the resulting initial-value problem by numerical integration as described in Materials and Methods. The dynamic behavior of this model is illustrated in **Figure 3C** which shows the result of a simulation, demonstrating that the repressor-production model is indeed capable of producing a pulse of gene expression that coincides with the period of flagellar growth.

### Model of repressor sequestered by flagellum

The fact that maximum expression levels are seen during the time of fastest flagellar growth (**Figure 1B**), suggested a second potential model in which the process of flagellar growth relieves gene repression. As with the previous model, we propose that flagellar gene expression is under control of a repressor. But in this case, the repressor is not generated by the flagellum, but instead is the protein product of a nuclear gene that auto-inhibits its own transcription and that is physically transported into the growing flagellum (**Figure 3B**). In this model, when the flagellum is detached, there is no immediate effect on transcription because the repressor is still present in the cytoplasm at its steady state level. As the flagellum begins to grow, repressor would be incorporated into the growing flagellum at a rate exceeding its rate of translation in the cytoplasm, resulting in a net loss of repressor from the cell body. This depletion of the repressor would then trigger transcription of flagellar genes including the repressor itself. As the flagellum reaches its equilibrium length and its growth rate slows, the rate of repressor depletion from cytoplasm would drop, allowing translation in the cell body to re-accumulate repressor in the cell body. Eventually the repressor level would increase enough to shut off further synthesis and the entire system would then be back at its original steady state. Unlike the first model, which clearly represents a negative feedback control on flagellar growth in that longer flagella make more repressor, in this case, faster flagellar growth would lead to more sequestration of repressor and therefore even more production of precursors to support further growth. The potential may thus exist for positive feedback to create an unstable “run-away growth” situation, in which upregulation of flagellar genes causes faster assembly, leading to further upregulation of assembly. Thus, stability of the model becomes a more serious concern.

Denoting as above the flagellar lengths, total pool of flagellar precursor protein, and message encoding precursor as L, M, and P, the model includes identical equations to the first model

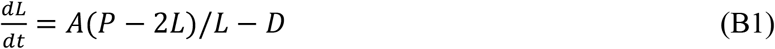

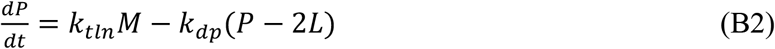

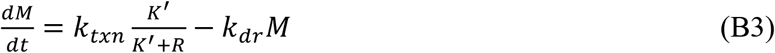

As in the previous model, R describes the concentration of repressor protein in the cell body. In this case, since R is transcribed and translated in the cell body, and transported into the flagellum along with the precursor proteins, we assume that at any point in time the concentration of repressor is proportional to the quantity of flagellar precursor protein in the cell body such that R=F(P-2L). In this case, K’ describes the dissociation constant for binding of the repressor not only to the promotor of genes encoding flagellar proteins but also to its own promotor.

It will prove convenient to focus on the cytoplasmic precursor pool Y rather than the total protein quantity P. To this end, we rewrite equation 1 as follows

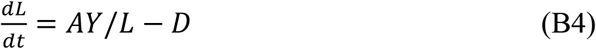

Next we replace equation B2 with a corresponding equation describing the dynamics of the cytoplasmic pool Y as follows

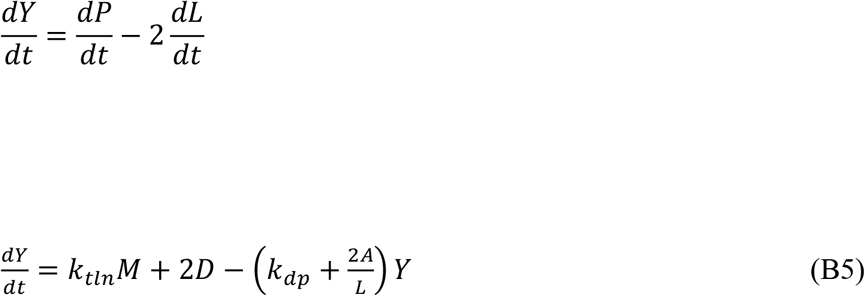

Making the substitution K=K’/F and replacing (P-2L) with the variable Y we obtain a simplified equation for mRNA dynamics

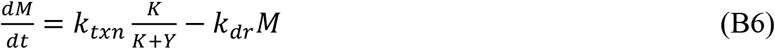

This model depends on seven parameters, which are summarized in Table 1. As derived in Materials and Methods, equations B4–B6 have a stead state solution:

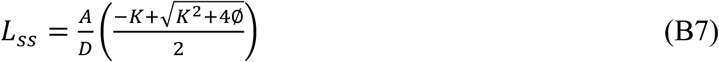

Where we have defined

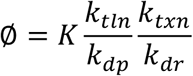

Thus a non-zero steady state solution for flagellar length always exists given the way our parameters are defined. Consistent with intuition, increasing the constant A describing IFT activity leads to increased flagellar length, as does decreasing the flagellar disassembly rate D. As derived in Materials and Methods, linear stability analysis indicates that this steady state solution is stable.

**Table 1.**
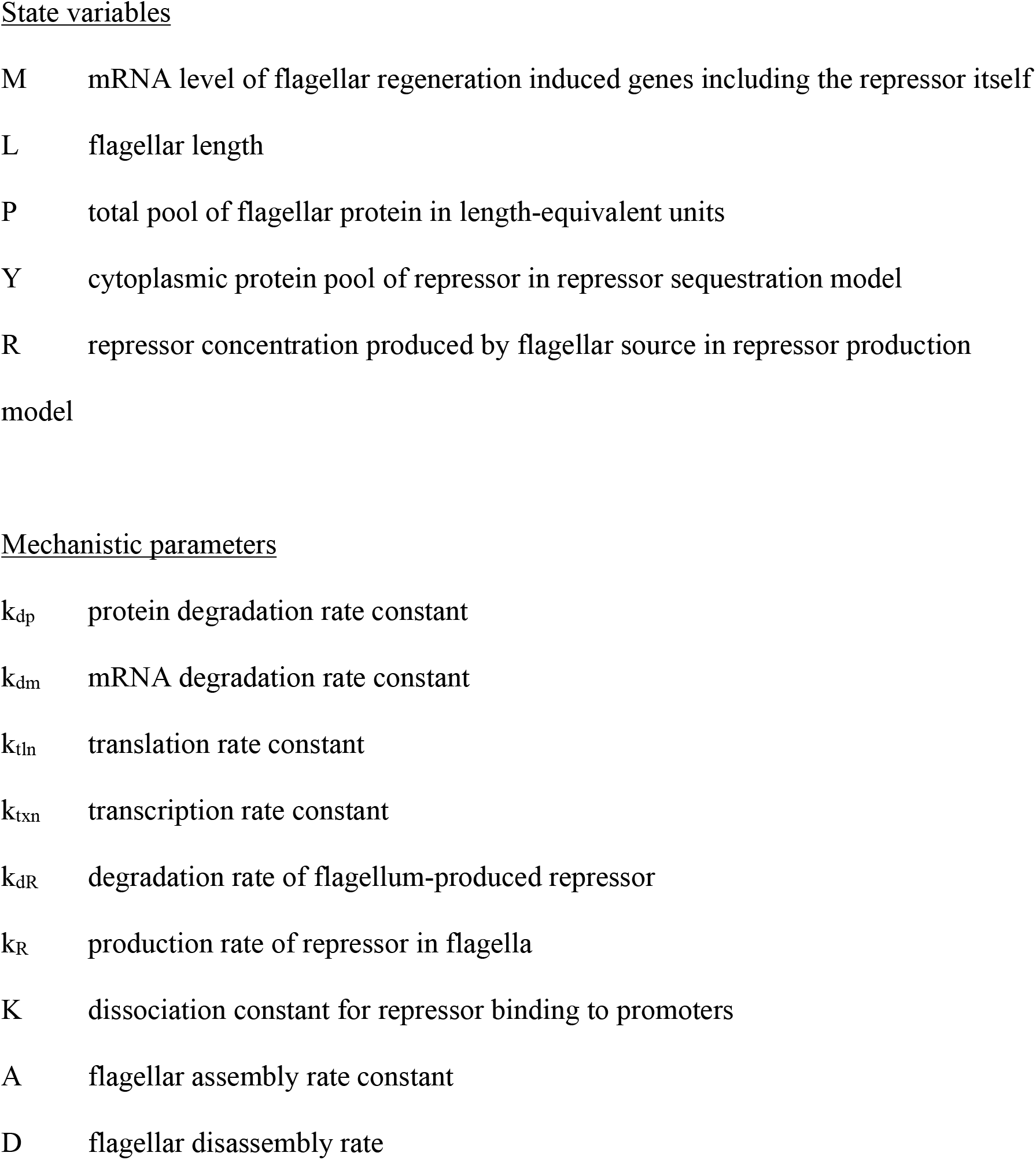
Parameters of repressor production and sequestration models.

State variables
|  |  |
| --- | --- |
| M | mRNA level of flagellar regeneration induced genes including the repressor itself |
| L | flagellar length |
| P | total pool of flagellar protein in length-equivalent units |
| Y | cytoplasmic protein pool of repressor in repressor sequestration model |
| R | repressor concentration produced by flagellar source in repressor production model |
| Mechanistic parameters |  |
| $k_{dp}$ | protein degradation rate constant |
| $k_{dm}$ | mRNA degradation rate constant |
| $k_{tln}$ | translation rate constant |
| $k_{txn}$ | transcription rate constant |
| $k_{dR}$ | degradation rate of flagellum-produced repressor |
| $k_R$ | production rate of repressor in flagella |
| K | dissociation constant for repressor binding to promoters |
| A | flagellar assembly rate constant |
| D | flagellar disassembly rate |

The dynamic behavior of the repressor sequestration model is illustrated in **Figure 3D** which shows that this model, like the repressor production model, is also capable of producing a pulse of gene expression that coincides with the period of flagellar growth. Consistent with the linear stability analysis, the system settles into a steady state in which the flagellar length does not change or increase out of control.

### Translation inhibition leads to increased gene expression

Extensive prior studies have shown that in the presence of translational inhibitors, flagella grow back to approximately 1/2 their initial length, making use of a pre-existing pool of precursor protein in the cytoplasm (Rosenbaum 1969). We asked whether the two models above might make testable predictions about transcriptional regulation in such cells.

In the case of the first model in which a repressor is produced within the flagellum at a rate proportional to flagellar length, the prediction is that since flagella never reach full length, the repressive signal will never reach its maximum level, leading to a sustained increase in transcription. Simulations of the model confirm this expectation (**Figure 3E**).

Within the repressor depletion model, the restoration of mRNA level back to baseline requires synthesis of new repressor protein to make up for the fraction of repressor that became incorporated into the flagellum. Consequently, if protein synthesis were prevented, transcription would initially occur at the same rate as in untreated cells, but then would not decrease because repressor never gets produced. Simulations of the model support this expectation (**Figure 3F**).

We conclude that both models, being based on negative regulation of transcription via mechanisms depending on protein synthesis (growth of flagella to full length in the first model, and production of a repressor protein in the second) make a similar prediction - that when protein synthesis is inhibited prior to flagellar regeneration, the result should be a sustained increase in transcript levels. This prediction stands in contrast to what one would expect if flagellar gene upregulation was based on production of a positive activator protein. In that case, inhibition of protein synthesis should prevent, rather than enhance, expression of flagella related genes.

When we measured mRNA levels during flagellar regeneration in cells treated with cycloheximide, we found (**Figure 3G,H**) that indeed, transcript levels were higher than in untreated cells at later timepoints, indicating that the pulsatile form of the transcription response was eliminated and replaced with a monotonic increase. Even at a very late time point, 140 min after the initial pH shock, (**Figure 3H**), transcript levels were substantially higher in cells treated with cycloheximide. The effect was not seen at earlier time points, nor was there any effect seen on non-flagellar control genes as in **Figure 3H**, indicating that cycloheximide does not simply increase mRNA levels uniformly. These results are therefore consistent with the predictions of the repressor-based models, but not with any model that requires synthesis of an activator protein to drive gene expression. Unfortunately, these experiments do not allow us to discriminate between the repressor production and repressor sequestration models, because both models make similar predictions. We also note that some of the increase in transcript abundance could be due to an effect of cycloheximide on the stability of flagella-related transcripts (Baker 1986).

### Mutants in the IFT pathway decrease gene induction

The repressor production model is based on production of a repressive signal by full length flagella, such that impairment of flagellar assembly, for example by inhibition of IFT, should lead to an increase in gene expression due to lack of the full repressive signal. The repressor sequestration model is based on import of a repressor into the flagellum to keep it away from the nucleus. Presumably, this sequestration would require intraflagellar transport, such that impairment of IFT and flagellar assembly should lead to a decrease in gene expression. As shown in **Figure 4A,B**, the two models make distinct predictions about the phenotype that should result from mutations that impair IFT. In the repressor production model, impairment of flagellar assembly leads to increased gene expression due to reduced production of the repressor (**Figure 4A**). In the repressor sequestration model, impairment of flagellar assembly leads to reduced gene expression, because the repressor is sequestered more slowly, allowing it to build up in the cytoplasm (**Figure 4B**). Experimental measurement of gene induction in IFT mutants with impaired flagellar assembly or transport could thus potentially help to distinguish these two models. We note that, as can be seen in equation (B7), the steady-state gene expression level in the repressor sequestration model is not predicted to be altered if IFT is altered (as reflected by the parameter A in the model). Rather, the effect should only be seen on the peak expression achieved during regeneration, as is seen in the simulation example of **Figure 4B**.

**Figure 4.**
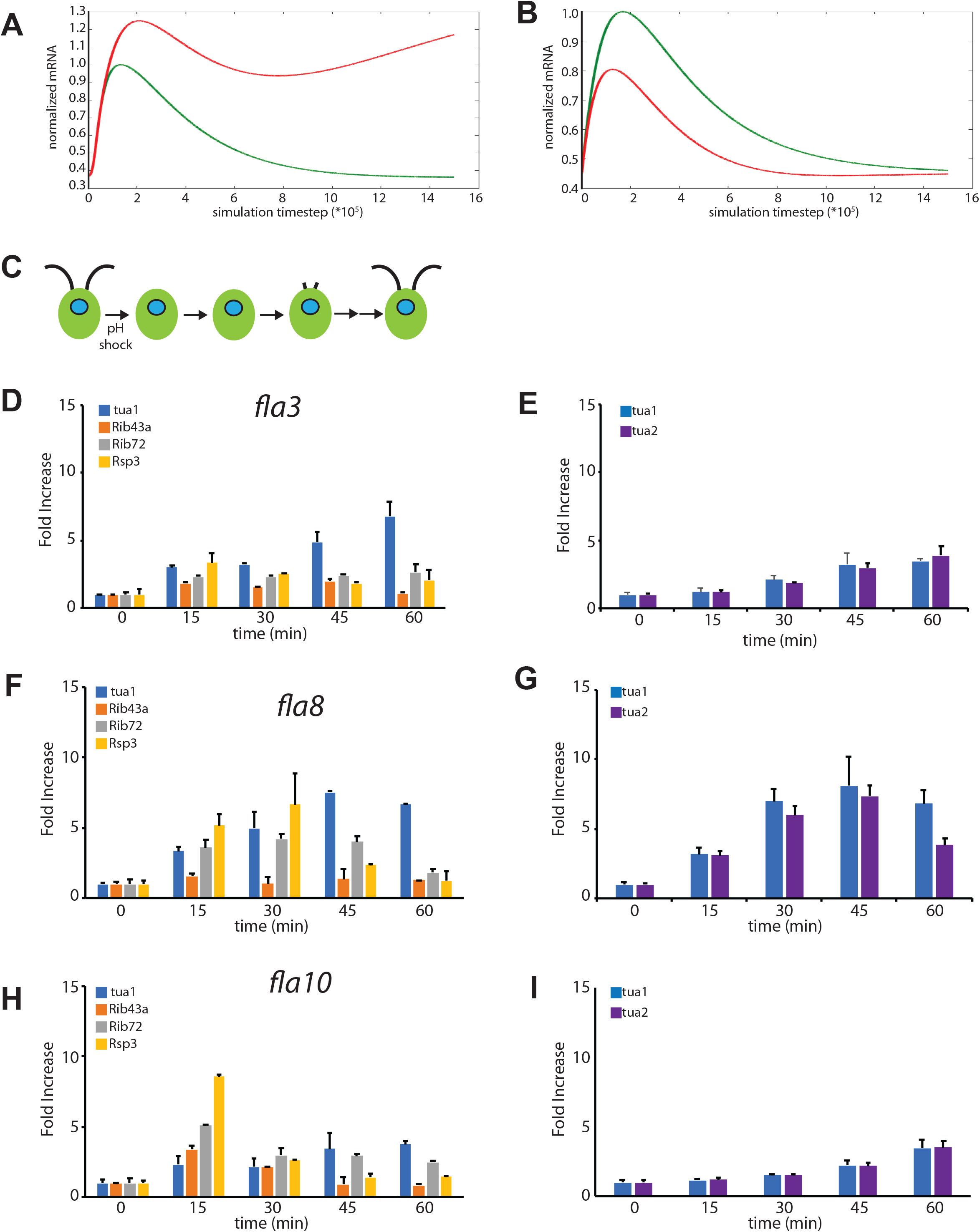
IFT mutants reduce upregulation of flagella related genes. (**A**) Prediction of the effect of reduced IFT in the repressor production model, in which a repressor is generated in the flagella proportional to flagellar length. Simulation shows mRNA levels using parameters from the plot in **Figure 3C** (green) with a second plot showing mRNA levels when the efficiency of IFT is reduced four-fold (red). Reduction in IFT leads to an increase in peak mRNA levels. (**B**) Prediction of the effect of reduced IFT in the repressor sequestration model, in which a repressor is generated in the cell body and then sequestered in flagella by IFT-mediated import. Simulation shows mRNA levels using parameters from the plot in **Figure 3D** (green) with a second plot showing mRNA levels when the efficiency of IFT is reduced four-fold (red). In contrast to the repressor production model, in the repressor sequestration model, reduction in IFT leads to a decrease in peak mRNA levels. **C**) Phenotype of conditional IFT mutants grown at the permissive temperature, as exemplified by the *fla3* mutant. Cells have normal length flagella, but show a prolonged delay in regeneration due to a defect in IFT. (**D**) Flagella-related gene expression following pH shock in *fla3* mutant quantified by qPCR. (**E**) Expression of tubulin genes following pH shock in *fla3* mutant. (**F**) Flagella-related gene expression following pH shock in *fla8* mutant quantified by qPCR. (**G**) Expression of tubulin genes following pH shock in *fla8* mutant. (**H**) Flagella-related gene expression following pH shock in *fla10* mutant quantified by qPCR. (**I**) Expression of tubulin genes following pH shock in *fla10* mutant.

In order to test the predictions of these two models, we require a mutation that will be defective in flagellar assembly due to an impairment of IFT, but in order to compare results with regeneration in wild-type cells, we require mutants that have full length flagella prior to pH shock. In fact, a number of mutants in IFT-related proteins have exactly this phenotype. We consider the *fla3* mutation, which affects one subunit of the IFT kinesin motor.

The FLA3 gene, which encodes the non-motor subunit of the kinesin-2 that drives intraflagellar transport (Mueller 2004), was originally identified via a conditional mutation that causes complete lack of flagella at the restrictive temperature (Adams 1982). However, at the permissive temperature (21C), the flagella in *fla3* mutants (**Figure 4C**) are close to wild-type length, but they fail to regenerate in the normal time frame following pH shock, requiring many hours to reach full length (Mueller 2004). This delay in regeneration is apparently due to the fact that the frequency of anterior IFT trains is reduced in these mutants even at 21C (Mueller 2004). As shown in **Figure 4D-E,** flagellar gene induction in conditional *fla3* mutant cells at 21C is in fact greatly reduced compared to wild-type cells. This result is consistent with the prediction of the repressor sequestration model but not the repressor production model (**Figure 4A vs. B**).

A similar reduction in anterior IFT frequency, accompanied by delayed regeneration of flagella, has also been reported for conditional *fla8* and *fla10* mutants grown at 21C (Iomini 2001). FLA8 and FLA10 encode the two motor subunits of heterotrimeric kinesin-2 (Walther 1994; Cole 1998; Miller 2005) and thus are expected to act at the same step of transport as FLA3. As shown in **Figure 4F-I**, we observed reduced peak levels of mRNA in both *fla8* and *fla10* mutants, similar to what we observed in the *fla3* mutant, and matching the prediction of the repressor sequestration model.

To further test this prediction, we measured expression in cells carrying the *fla1* mutation (originally named dd-a-6), in which flagella at 21C are full length but fail completely to regenerate (Huang 1977). This mutation is actually an allele of the FLA8 gene (Miller 2005) and thus affects flagellar assembly via a defect in IFT. If expression was triggered by a positive detector of flagellar loss, or inhibited by a repressor produced by the assembled flagellum, gene expression should be high in such a mutant. Instead, we find that tubulin expression in this mutant is severely reduced to levels close to those seen in the bld mutants (**Supplemental Figure S1B**), again consistent with the repressor sequestration model.

### Microtubule polymerization inhibitors decrease late but not early gene induction

The reduced expression in IFT mutants could indicate a direct role of IFT in sequestering the repressor inside the flagellar compartment. But it could also reflect an indirect effect, where the repressor is sequestered anchoring the repressor protein onto the axoneme, trapping it inside the flagellum and preventing it from reaching the nucleus. In such a version of the model, the effect of IFT mutants seen in **Figure 4** would be an indirect one, reflecting the requirement of IFT to build the axoneme onto which the repressor is anchored. To distinguish between these possibilities, we tested whether oryzalin, a specific chemical inhibitors of algal microtubule polymerization (5James 1988), and which blocks flagellar assembly to an extent similar to or greater than the conditional mutants in IFT, would similarly reduce gene expression levels. As seen in **Figure 5**, oryzalin treatment had very little effect on gene expression levels, although a modest reduction was seen a later timepoints. These results suggest that impaired axonemal assembly has a far less significant effect on flagella-related gene expression than impairment of IFT.

**Figure 5.**
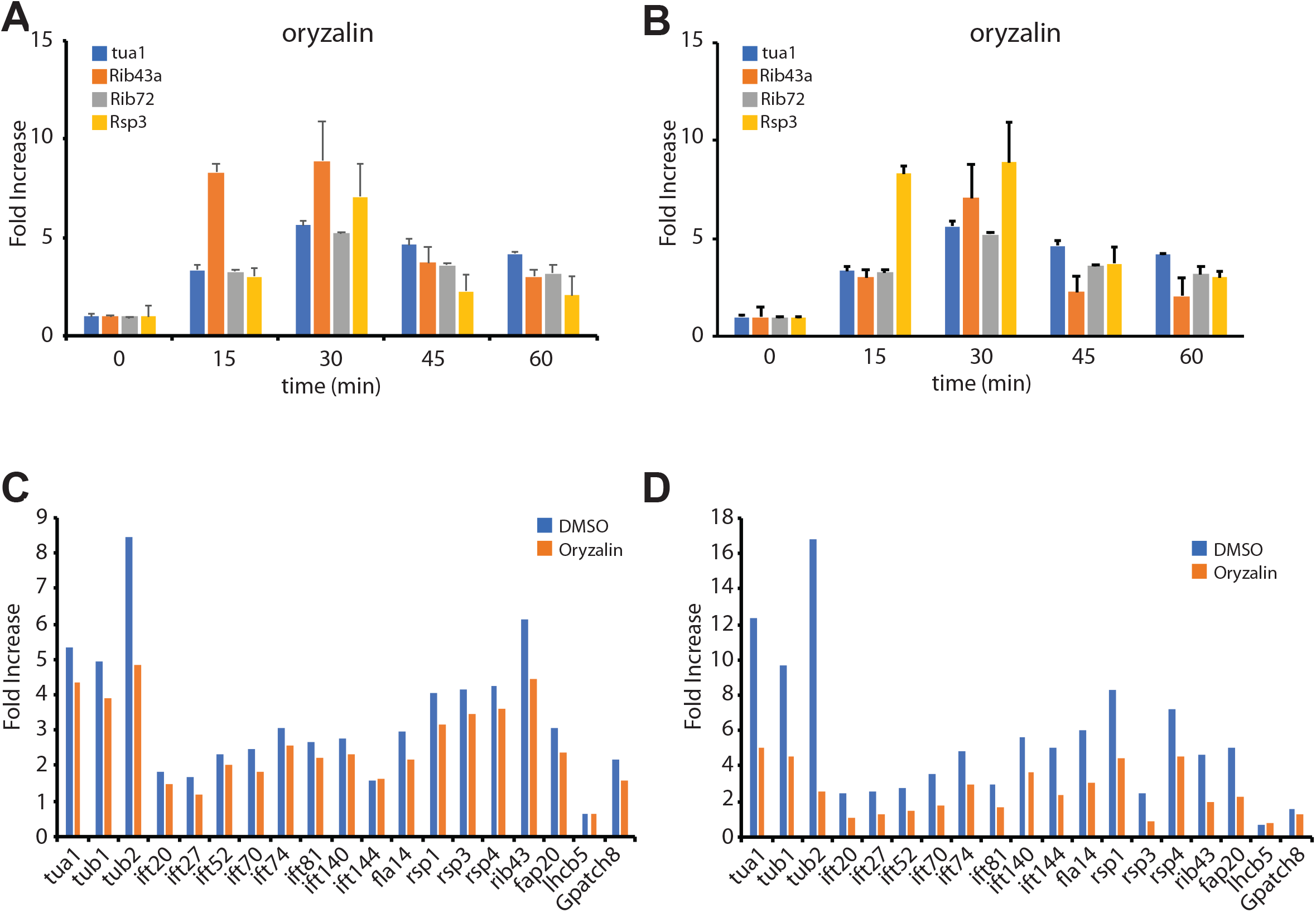
Effect of microtubules inhibitors on flagella related gene expression after pH shock. (**A,B**) Effect of 3.5 μM oryzalin on gene expression judged by qPCR. Graphs represent two independent experiments. (**C,D**) Effect of 3.5 μM oryzalin on gene expression after pH shock quantified by Nanostring detection. Panels C and D represent expression 30min and 45 min after deflagellation, respectively.

## Discussion

### Regulation of transcription by a negative regulator - comparison to other studies

Neither the pH shock itself, nor the subsequent loss of flagella, are sufficient to trigger the upregulation of flagellar genes. This conclusion is based on the fact that pH shock does not drive a transcriptional burst in cells that lack flagella to start with (**Figure 2**), and that in mutants in IFT proteins that start out with full length flagella, and lose them normally during pH shock, but cannot regenerate them at normal rates, gene induction is greatly reduced (**Figure 4**). The fact that expression is reduced, rather than enhanced, in IFT mutants with slowly regenerating flagella suggests that a negative regulator may be sequestered in the flagellum rather than produced by the flagellum, and the fact that IFT mutants have much more severe effect than inhibition of microtubule polymerization suggests that the sequestration may be directly carried out by IFT mediated import.

A study in which flagellar regeneration was inhibited by depletion of calcium (Cheshire 1991) led to conclusions highly consistent with our model. In that study, when cells were deflagellated by pH shock in media from which the calcium had been removed by chelators, flagella did not regenerate, and flagellar gene induction did not take place. If, in such cells, calcium was restored to the media two hours after deflagellation, flagella began to assemble, and at that point a pulse of flagellar gene expression was observed. That study argued that flagellar assembly was sufficient to induce gene expression.

According to our repressor-sequestration model, any perturbation leading to increased IFT should cause a pulse of gene expression. It has been shown that treatment of *Chlamydomonas* cells with Li^+^, known to stimulate flagellar growth, leads to upregulation of flagella-related genes (Periz 2007). We previously showed that Li^+^ causes an increase in IFT injection rates (Ludington 2013) thus the observed induction of flagellar genes when lithium is added may thus be consistent with the idea that IFT is involved in sequestering a repressor in the flagellum. The same study (Periz 2007) also found that when cycloheximide was added in addition to Li^+^, a larger increase in flagella related mRNA was observed, which is consistent with our observations during regeneration. They also found that cycloheximide by itself, in the absence of Li^+^, was not sufficient to cause an increase in abundance of flagella related mRNA.

Conversely, the observation that shortening of flagella induced by IBMX causes a decrease in flagella-related gene expression (Lefebvre 1980; Chamberlain 2008) is also be consistent with the model because as flagella disassemble, their contents return to the cell body, and this would include the putative repressor. In contrast, with a repressor-production model, disassembly of the flagellum should lead to less repressor, and more gene expression.

Larkin et al. (1989) described a mutant that regenerated flagella slowly at the restrictive temperature, but in which tubulin gene induction was normal. This phenotype was thus more reminiscent of the effects of tubulin inhibitors. Regrettably, the identity of the gene mutated in that study is not known, but we speculate that it might indicate a gene that normally plays a role in axonemal assembly rather than IFT, such that the mutant would still be able to sequester a repressor normally.

The fact that gene expression is only slightly impaired when flagellar regeneration is prevented with microtubule inhibitors (**Figure 5**) is largely consistent with published results. Rosenbaum et al (1969) showed that the functional pool of flagellar precursor increased over time when cells were deflagellated in presence of colchicine, indicating that genes are induced. It was further reported that tubulin protein synthesis occurs following deflagellation in the presence of the microtubule inhibitor colchicine (Lefebvre et al., 1978). Weeks (1977) found that tubulin genes were still expressed when cells were deflagellated after treatment with IBMX to prevent axonemal regrowth. On the other hand Minami (1981) found that when regeneration was blocked with a different microtubule inhibitor, APM, the cells did not induce tubulin gene expression. The autoregulation of tubulin at a post-transcriptional level may complicate interpretation of experiments with tubulin inhibitors.

Our data presented here with oryzalin, along with the prior data just discussed, indicate that axonemal assembly is not required to sequester a repressor in the flagellum. But where is the repressor being sequestered, if axonemes are not required? Electron microscopy studies have shown that when flagellar regeneration in blocked with microtubule inhibitors, *Chlamydomonas* cells still form a short empty flagellar stump consisting of a flagellar membrane devoid of an axonemal structure (Jarvik and Chojnacki 1985). We propose that IFT is used to import the repressor into this flagellar stump. This possibility will require further experimental investigation but, if true, it would potentially provide a strategy for identifying the repressor molecule. An alternative possibility is that the IFT kinesin-2 complex plays a more direct role in blocking or inhibiting a repressor gene expression, in a manner independent of its role in flagellar assembly. Our present data cannot rule out such a possibility.

### Possible molecular identity of the negative regulator

Our model suggests that flagella related gene induction can be explained in terms of a negative regulator that is incorporated into the growing flagellum, and that shuts down flagella-specific gene expression when it accumulates in the cell body. Our model does not make any direct assumption about the molecular identity of this regulator. One simple possibility could be a transcriptional inhibitor that shuttles between the nucleus and the flagellum. Such a scheme, in which a transcription regulator can move from an organelle to the nucleus, is not without precedent. For example the famous SREBP protein resides in the endoplasmic reticulum when cholesterol is abundant but part of the protein relocates to the nucleus to active expression of genes involved in cholesterol synthesis when cholesterol levels drop. We previously analyzed a small set of insertional mutants in flagellar proteins containing putative DNA binding domains, but saw no effect on flagellar length or assembly (Perlaza 2021). However, that study was limited to DNA binding proteins for which insertional mutants already existed in the CLiP mutant library collection (Li 2016), leaving open the possibility that one or more other flagellar proteins with DNA binding domains could act as the unknown repressor.

However, it is equally possible that the negative regulator sequestered by IFT is not itself a transcriptional repressor, but instead acts more indirectly. For example, it could be a kinase or a protease that targets a transcriptional activator to inactivate or degrade it, or part of a more extensive signal transduction pathway. In fact, the actual molecule sequestered inside the flagellum could be anything upstream of the final DNA binding transcriptional regulator. Given that there are so many possible ways to construct a negative regulatory pathway, we suggest that an unbiased genetic approach may be more productive. The repressor sequestration model provides some potential considerations for designing such a screen.

First, if control of flagellar gene expression is mediated by a negative regulator, then loss of function mutations in the repressor would be expected to cause constitutive gene expression. Previous attempts to screen for mutants affecting flagella specific gene expression were designed to identify mutants that failed to show gene upregulation upon pH shock. If our model is correct, then it may prove far more productive to look for mutants that fail to turn off gene expression after flagella have finished assembling.

Second, because repression only occurs when flagella regrow, and not simply as a result of pH shock on cells lacking flagella (**Figure 2**), it will only be informative to consider mutants that have normal length flagella prior to pH shock. For example, we previously found that a mutant strain carrying the short-flagella mutation *shf2* failed to upregulate flagella-related genes following pH shock (Kannegaard 2014). Although *shf2* mutant cells had previously been reported to have flagella, albeit short ones (Kuchka and Jarvik 1978), at least in our *shf2* lab strain, flagella were absent from most cells, even pre-shock. Hence the lack of induction is consistent with the requirement for flagellar assembly and does not provide any evidence that the SHF2 gene product plays any role in transcriptional regulation. Similarly, a mutant was reported in a transcription factor XAP5 that resulted in loss of flagellar gene up-regulation (Li 2018), but these mutant cells lacked flagella, hence the effect on transcription is likely to be indirect.

Taking these considerations into account, the ideal screen would look for mutants in which cells have flagella, but constitutively express flagella related genes. The power of *Chlamydomonas* genetics should make such a screen possible, when combined with high throughput assays for expression.

### Implications for biological feedback control systems

What are the implications of the repressor sequestration model from a control system perspective? Most existing models for organelle size regulation are essentially proportional controllers, in the sense that they invoke a mechanism whereby the assembly process of the organelle is regulated as a function of the current size of the organelle relative to some target size. However, human designed feedback controllers often include not just proportional control but also integral control, which depends on the time-integral of the error, and derivative control, which depends on the rate of change of the error. Combination of all three types of feedback results in a PID controller, which is standard in industrial control applications. There has been recent interest in building synthetic PID controllers in cells (Chevalier 2019), but it remains an interesting open question to determine to what extent do cells naturally employ these types of control schemes.

The fact that mRNA levels of flagellar genes are proportional to the rate of flagellar growth **Figure 1C** leads us to speculate that the transcriptional control of flagellar gene expression might represent a form of derivative feedback. In this case the feedback is positive - the faster the flagellum grows, the more repressor is sequestered, and the higher the rate of flagellar precursor production, ultimately leading to more growth. Positive derivative feedback is not necessarily a bad thing as long as the system is able to reach its target point and remain there stably. Indeed, the gain for the derivative term in a PID controller is sometimes deliberately made opposite the gain for the proportional and integral terms, in order to adjust the speed of system response. In the present case, the fact that production of precursor turns on when flagella are in their initial rapid growth phase may help flagella reach steady state length more rapidly. This idea is consistent with the fact that initial growth of flagella uses pre-existing precursor pools, and newly synthesized precursors only become necessary in the later stages of assembly to reach final length (Rosenbaum 1969).

## Materials and Methods

### Quantitative measurement of mRNA levels in Chlamydomonas

Cells were grown in 10mL of TAP media (Harris, 1989) in a test tube on a roller drum at 21C for 2 days, then transferred 5mL of starter culture into a 250mL flask which contained 95mL of TAP. We placed the flask into the incubator at 21C and allowed it to bubble for two days. For experiments inhibiting microtubule polymerization, we added 3.5μM oryzalin 15 minutes before deflagellation and conducted the experiment in the continuous presence of the inhibitor.

Deflagellation was accomplished by pH shock. We first aliquot 25mL of cells into a 50mL conical tube, we add 1.25mL of 0.5N Acetic Acid, allowing the cells to mix by inversion for 1 minute before bringing the pH back to normal by adding 1mL of 0.5N Sodium Hydroxide and inverting for 1 minute before placing back in the 21C incubator and bubbling. Samples were collected at the indicated timepoints, mixed with Trizol (Invitrogen, Carlsbad, California, United States) and stored at −80. RNA was prepared by chloroform extraction followed by isopropanol precipitation. Concentration was determined using Nanodrop (ND1000, NanoDrop) and then stored at −80 C.

Samples were DNAse treated then used for RT-PCR using 500ng of Random Primers (Invitrogen, Carlsbad, California, United States) and the Super Script II kit(Invitrogen, Carlsbad, California, United States) following manufacturer’s instructions with the addition of Rnasin (Promega). Concentrations were checked by Nanodrop.

Quantitative PCR (qPCR) was performed using 5uL of the total cDNA (derived from 100ng RNA) per reaction with 20uL of cocktail containing 2x SYBR green supermix (Biorad, Hercules, California, United States), gene-specific Primer Pairs (IDT, Iowa, United States), Betaine (Sigma-Aldrich Inc, St. Louis, Missouri, United States), and water (Sigma-Aldrich Inc, St. Louis, Missouri, United States).

Primers for qPCR were as follows:

RBCS2

L 5’ ACAAGGCCTACGTGTCCAAC 3’
R 5’ ATCTGCACCTGCTTCTGGTT 3’

Rsp3

L 5’ GCCATCACCCAGATCGAG 3’
R 5’ CCTCCTCCTCCAGCACCT 3’

Rib72

L 5’ TTCATGATGGACGACTCCAA 3’
R 5’ GCGGCTTGTAAATCTTCTGG 3’

Rib43a

L 5’ CGCAGATGGAGGAGAAGAAG 3’
R 5’ CCTCTTGCTGCTTTTTGAGG 3’

Tua2

L 5’ GCCAATAGAGGCACGGTCGTGGA 3’
R 5’ GGCGTGATCTGAGGCTTCGTTGG 3’

For Nanostring quantification, cells were lysed in trizol and RNA collected using the Direct-zol RNA miniprep method (Zymo Research) and analyzed by Nanostring detection using the probeset detailed in **Supplemental Table S1** and nCounter Sprint Cartridge. Data was analyzed using the nCounter software package.

For measuring flagellar length (Figure 1B,C), cells from the same sample as used for mRNA quantification were fixed in 1% glutaraldehyde. Cells were imaged on a Deltavision microscopy system using a 60x air lens using DIC optics with an air condenser. Three dimensional datasets were collected with a z-spacing of 0.2 microns. Flagella were traced in three dimensions to calculate length. For calculation of flagellar growth rate (Figure 1C), slopes of the best fit line were fit in a sliding window of five timepoints centered on each point, except for the first four timepoints which were based on correspondingly fewer points.

### Analytical solution of models

Each of the two models (repressor production, equations A1–A4, and repressor sequestration, equations B4–B6) were solved analytically for a steady state solution, which was then tested for local stability by linear stability analysis. Numerical exploration of the models will be addressed in a subsequent section below.

#### Steady state solution of repressor production model

We solve for the steady state solution of equations A1–A4 by setting each rate to zero, from which we obtain:

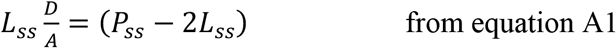

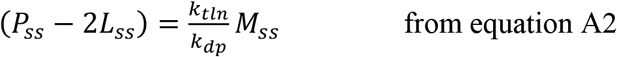

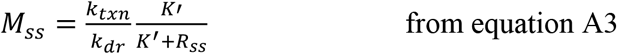

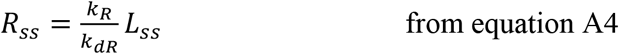

Defining γ=k_R_ /(K’ k_dR_)

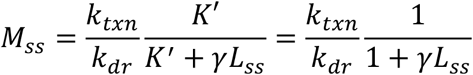

Hence

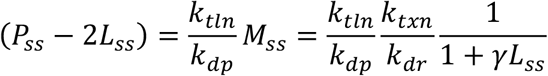

Defining

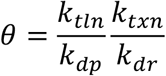

We get

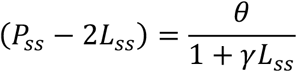

Substituting (P-2L) into the first equation, we obtain

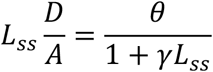

Which we solve as a quadratic equation to yield the steady state length:

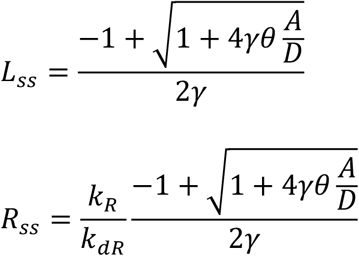

#### Steady-state solution of repressor sequestration model

Next we find the steady state solution to the system of equations B4–B6 describing the repressor sequestration model.

Setting dL/dt=0 in equation B4 we obtain

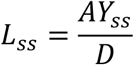

Setting dM/dt = 0 in equation B6 we obtain

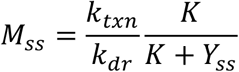

Finally setting dY/dt =0 in equation B5 we obtain

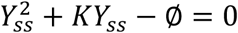

Where we have defined

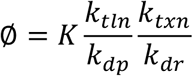

We then obtain

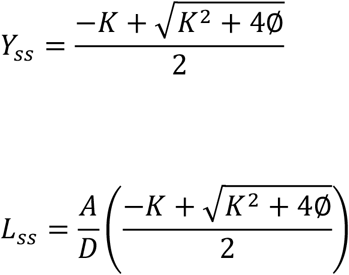

This model has seven parameters, which are summarized in Table 1.

#### Stability analysis of repressor-sequestration model

To determine if the steady state solution is a stable fixed point, we linearize the system (B4-B6) in the vicinity of the fixed point. Representing the state of the system with the vector [M, Y, L], the Jacobian matrix at the fixed point is:

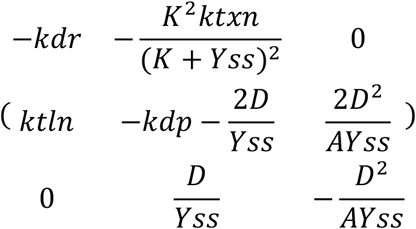

Which has the characteristic polynomial

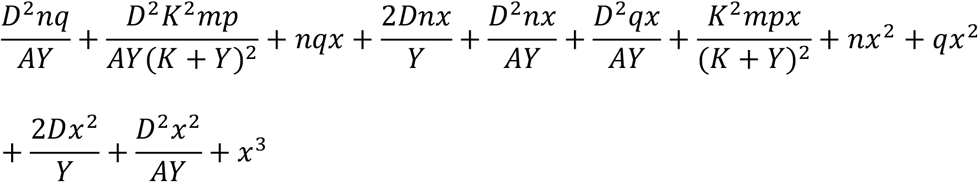

Where we have replaced rate constant names k_txn_, k_dr_, k_tln_, and k_dp_ with the letters m,n,p, and q respectively. According to the Routh-Hurwitz criterion, all roots of a cubic polynomial have strictly negative real parts if and only if the three coefficients ao,ai, and a2 corresponding to the 0, 1, and 2 powers of x, are all positive and *a*_2_ * *a*_1_ > *a*_0_. Given that all parameters of the model are strictly positive, and that the steady state value of Y which appears in the Jacobian is also strictly positive, it is clear that all coefficients of the polynomial are positive. The criterion a2*a1>a0 is guaranteed to be satisfied because for each of the two terms composing

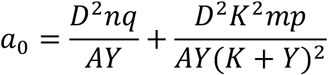

It can be seen that there is a corresponding term in the product a1*a2. Since every term in a_1_ and in a_2_ are strictly positive, the additional terms in their product are positive, hence it will always be the case that a1*a2>0 and the Routh-Hurwitz criterion is satisfied. Thus, all eigenvalues of the Jacobian for the linearized system have negative real parts, so the fixed point is stable for the linearized system. Moreover, this result also establishes that the fixed point is hyperbolic, hence the Hartman-Grobman theorem indicates the stability properties of the linearized system will carry over to the nonlinear system at least in a sufficiently small neighborhood around the fixed point.

### Numerical simulations of repressor production and sequestration models

The preceding analysis lets us make statements about the steady state solution and the stability of that solution in a local region around the fixed point, but does not tell us anything about the dynamics of how the system evolves over time starting from an initial condition such as that seen in flagellar regeneration. We implemented numerical simulations of the systems of equations for the two models discussed above using the Euler method with 1500000 steps. The purpose of these simulations was just to determine the qualitative behavior of the systems, and not to make detailed numerical predictions. Hence, the following simple choices of parameters were used.

repressor production model:

|  |  |  |
| --- | --- | --- |
| $k_{dp}$ | protein degradation rate constant | 0.05 |
| $k_{dr}$ | mRNA degradation rate constant | 0.05 |
| $k_{tln}$ | translation rate constant | 1 |
| $k_{\text{txn}}$ | transcription rate constant | 1 |
| $k_{\text{dR}}$ | degradation rate of flagellum-produced repressor | 1 |
| $k_{\text{R}}$ | production rate of repressor in flagella | 5 |
| $K$ | dissociation constant for repressor binding to promoters | 2 |
| $A$ | flagellar assembly rate constant | 2 |
| $D$ | flagellar disassembly rate | 1 |

repressor sequestration model:

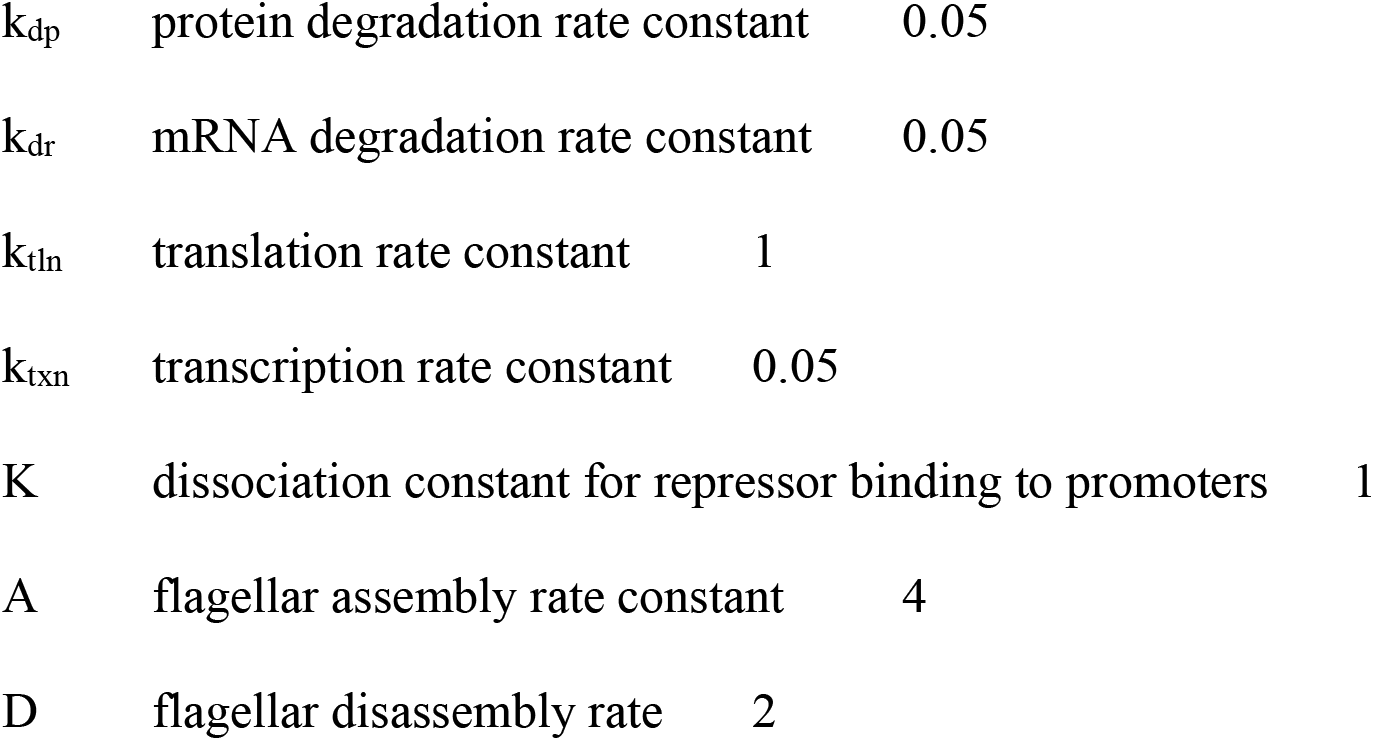

|  |  |  |
| --- | --- | --- |
| $k_{\text{dp}}$ | protein degradation rate constant | 0.05 |
| $k_{\text{dr}}$ | mRNA degradation rate constant | 0.05 |
| $k_{\text{tln}}$ | translation rate constant | 1 |
| $k_{\text{txn}}$ | transcription rate constant | 0.05 |
| $K$ | dissociation constant for repressor binding to promoters | 1 |
| $A$ | flagellar assembly rate constant | 4 |
| $D$ | flagellar disassembly rate | 2 |

The simulations were initialized to the steady state value of the state variables, calculated using the analytical solutions described above. After this initialization phase, flagellar regeneration was simulated by setting L to 0.1, and reducing the total protein pool (P in the repressor production model, and Y in the repressor sequestration model) by twice the pre-shock steady state flagellar length, in order to represent the protein lost during flagellar shedding.

To simulate effects of IFT mutants, the parameter A was reduced by a factor of 4. Results were plotted by normalizing to the steady state mRNA level using the original value of A, to reflect the fact that the fla mutants, being grown at permissive temperature, started out with full length flagella. To simulate the effect of cycloheximide, the value of k_tln_ was set to zero, and the simulation re-run.

The simulation programs were implemented in Matlab are available on github at the following link: https://github.com/WallaceMarshallUCSF/flagella-repressor-models

## ACKNOWLEDGMENTS

We thank Elisa Kannegaard for advice about qPCR and members of the Marshall lab for helpful discussions. This work was supported by NIH grant R35 GM130327.

---

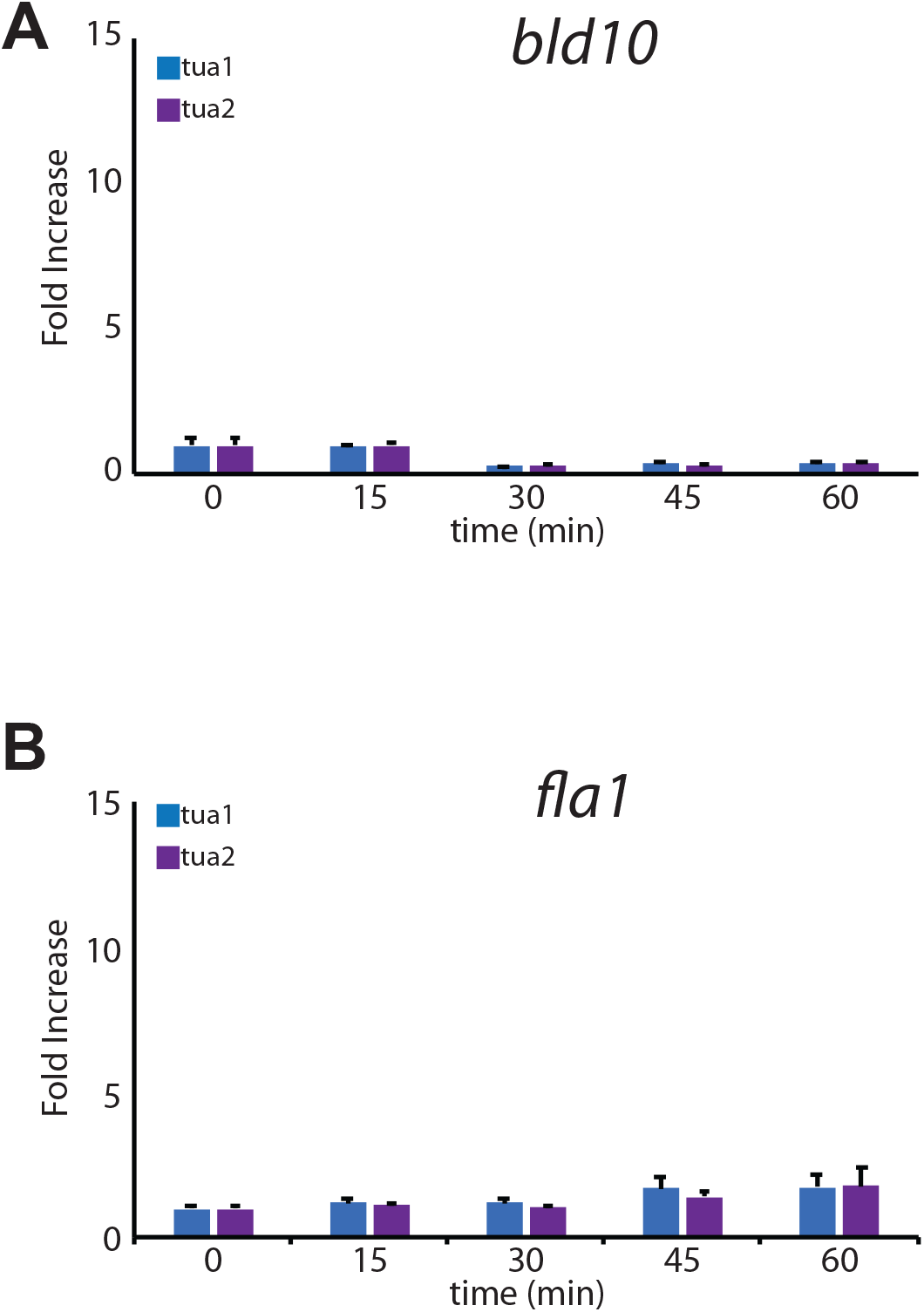

## References

Adams GMW, Huang B, Luck DJ (1982). Temperature-sensitive assembly-defective flagella mutants of *Chlamydomonas reinhardtii*. Genetics 100, 579–586

Albee AJ, Kwan AL, Lin H, Granas D, Stormo GD, Dutcher SK. (2013). Identification of cilia genes that affect cell-cycle progression using whole-genome transcriptome analysis in Chlamydomonas reinhardtii. G3 (Bethesda). 3, 979–91.

Baker EJ, Schloss JA, Rosenbaum JL. (1984). Rapid changes in tubulin RNA synthesis and stability induced by deflagellation in Chlamydomonas. J Cell Biol 99, 2074–2081

Baker EJ, Keller LR, Schloss JA, Rosenbaum JL. (1986). Protein synthesis is required for rapid degradation of tubulin mRNA and other deflagellation-induced RNAs in *Chlamydomonas reinhardtii*. Mol. Cell. Biol. 6:54–61.

Bhogaraju S, Cajanek L, Fort C, Blisnick T, Weber K, Taschner M, Mizuno N, Lamla S, Bastin P, Nigg EA, Lorentzen E. (2013). Molecular basis of tubulin transport within the cilium by IFT74 and IFT81. Science. 341, 1009–12

Bhogaraju S, Weber K, Engel BD, Lechtreck KF, Lorentzen E. (2014). Getting tubulin to the tip of the cilium: one IFT train, many different tubulin cargo-binding sites? Bioessays 36, 463–7

Brazelton WJ, Amundsen CD, Silflow CD, Lefebvre PA. (2001). The bld1 mutation identifies the Chlamydomonas osm-6 homolog as a gene required for flagellar assembly. Curr Biol. 11, 1591–4.

Chamberlain KL, Miller SH, Keller LR. (2008). Gene expression profiling of flagellar disassembly in Chlamydomonas reinhardtii. Genetics. 179, 7–19

Cheshire JL, Keller LR. (1991). Uncoupling of Chlamydomonas flagellar gene expression and outgrowth from flagellar excision by manipulation of Ca2+ J. Cell Biol. 115, 1651–9.

Chevalier M, Gómez-Schiavon M, Ng AH, El-Samad H. (2019). Design and Analysis of a Proportional-Integral-Derivative Controller with Biological Molecules. Cell Syst. 9, 338–353

Cole DG, Diener DR, Himelblau AL, Beech PL, Fuster JC, Rosenbaum JL. 1998. Chlamydomonas kinesin-II-dependent intraflagellar transport (IFT): IFT particles contain proteins required for ciliary assembly in Caenorhabditis elegans sensory neurons. J Cell Biol. 141, 993–1008

Craft JM, Harris JA, Hyman S, Kner P, Lechtreck KF. (2015). Tubulin transport by IFT is upregulated during ciliary growth by a cilium-autonomous mechanism. J Cell Biol 208, 223–237.

Davies JP, Grossman AR. (1994). Sequences controlling transcription of the Chlamydomonas reinhardtii beta 2-tubulin gene after deflagellation and during the cell cycle. Mol. Cell Biol. 14, 5165–74.

Engel BD, Ludington WB, Marshall WF. (2009). Intraflagellar transport particle size scales inversely with flagellar length: revisiting the balance-point length control model. J. Cell Biol. 187, 81–9.

Evans JH, Keller LR. (1997). Calcium influx signals normal flagellar RNA induction following acid shock of Chlamydomonas reinhardtii. Plant. Mol. Biol. 33, 467–81.

Goehring NW, Hyman AA. (2012). Organelle growth control through limiting pools of cytoplasmic components. Curr. Biol. 22, R330–339.

Hao L, Thein M, Brust-Mascher I, Civelekoglu-Scholey G, Lu Y, Acar S, Prevo B, Shaham S, Scholey JM. (2011). Intraflagellar transport delivers tubulin isotypes to sensory cilium middle and distal segments. Nat. Cell Biol. 13, 790–8.

Hendel NL, Thomson M, Marshall WF. (2018). Diffusion as a Ruler: Modeling Kinesin Diffusion as a Length Sensor for Intraflagellar Transport. Biophys J 114, 663–674

Huang B, Rifkin MR, Luck DJ. (1977) Temperature-sensitive mutations affecting flagellar assembly and function in Chlamydomonas reinhardtii. J Cell Biol 72, 67–85

Iomini C, Babaev-Khainov V, Sassaroli M, Piperno G. (2001). Protein particles in *Chlamydomonas* flagella undergo a transport cycle consisting of four phases. J. Cell Biol. 153, 13–24.

Ishikawa H, Marshall WF. (2017). Testing the time-of-flight model for flagellar length sensing. Mol Biol Cell 28, 3447–3456

Ishikawa H, Moore J, Diener DR, Delling M, Marshall WF. (2022). Testing the ion-current model for flagellar length sensing and IFT regulation. bioRxiv 2022.08.20.504661

James SW, Ranum LP, Silflow CD, Lefebvre PA. (1988). Mutants resistant to anti-microtubule herbicides map to a locus on the uni linkage group in Chlamydomonas reinhardtii. Genetics 118, 141–7.

Jarvik JW, Chojnacki B. (1985). Flagellar morphology in stumpy-flagella mutants of Chlamydomonas reinhardtii. J. Protozool. 32, 649–56.

Kannegaard, E., E.H. Rego, S. Schuck, J.L. Feldman, and W.F. Marshall. (2014) Quantitative analysis and modeling of katanin function in flagellar length control. Mol Biol Cell. 25, 3686–3698.

Keller LR, Schloss JA, Silflow CD, Rosenbaum JL. (1984). Transcription of alpha-and beta-tubulin genes in vitro in isolated Chlamydomonas reinharti nuclei. J. Cell Biol. 98, 1138–1143.

Kozminski KG, Johnson KA, Forscher P, Rosenbaum JL. (1993). A motility in the eukaryotic flagellum unrelated to flagellar beating. Proc. Natl. Acad. Sci. USA 90, 5519–5523.

Kuchka MR, Jarvik JW. (1987). Short-flagella mutants of *Chlamydomonas*. Genetics 115, 685–691.

Larkin JC, Lefebvre PA, Silflow CD. (1989). A gene essential for viability and flagellar regeneration maps to the uni linkage group of Chlamydomonas reinhardtii. Curr. Genet. 15, 377–84

Lefebvre PA, Nordstrom SA, Moulder JE, Rosenbaum JL. (1978). Flagellar elongation and shortening in Chlamydomonas. IV. Effects of flagellar detachment, regeneration, and resorption on the induction of flagellar protein synthesis. J. Cell Biol. 78, 8–27.

Lefebvre PA, Silflow CD, Wieben ED, Rosenbaum JL. (1980). Increased levels of mRNAs for tubulin and other flagellar proteins after amputation or shortening of Chlamydomonas flagella. Cell 20, 469–77.

Lefebvre PA, Rosenbaum JL. (1986). Regulation of the synthesis and assembly of ciliary and flagellar proteins during regeneration. Ann. Rev. Cell Biol. 2, 517–46.

Li, X, Zhang, R, Patena, W, Gang, SS, Blum, SR, Ivanova, N, Yue, R, Robertson, JM, Lefebvre, PA, Fitz-Gibbon, ST, Grossman, AR, Jonikas, MC (2016). An Indexed, Mapped Mutant Library Enables Reverse Genetics Studies of Biological Processes in Chlamydomonas reinhardtii. Plant Cell 28, 367–387.

Li L, Tian G, Peng H, Meng D, Wang L, Hu X, Tian C, He M, Zhou J, Chen L, Fu C, Zhang W, Hu Z. (2018). New class of transcription factors controls flagellar assembly by recruiting RNA polymerase II in Chlamydomonas. Proc. Natl. Acad. Sci. U.S.A. 115, 4435–4440.

Lin H, Dutcher SK. (2015). Genetic and genomic approaches to identify genes involved in flagellar assembly in Chlamydomonas reinhardtii. Methods Cell Biol. 127, 349–86

Ludington WB, Wemmer KA, Lechtreck KF, Witman GB, Marshall WF. (2013). Avalanche-like behavior in ciliary import. Proc Natl Acad Sci US A 110, 3925–3930.

Ludington WB, Ishikawa H, Serebrenik YV, Ritter A, Hernandez-Lopez RA, Gunzenhauser J, Kannegaard E, Marshall WF. (2015). A systematic comparison of mathematical models for inherent measurement of ciliary length: how a cell can measure length and volume. Biophys J 108, 1361–1379

Marshall WF, Qin H, Rodrigo Brenni M, Rosenbaum JL. (2005). Flagellar length control system: testing a simple model based on intraflagellar transport and turnover. Mol. Biol. Cell 16, 270–278.

Marshall WF, Rosenbaum JL. (2001). Intraflagellar transport balances continuous turnover of outer doublet microtubules: implications for flagellar length control. J. Cell Biol. 155, 405–14.

Marshall WF. (2016). Cell geometry: how cells count and measure size. Ann. Rev. Biophys. 45, 49–64.

Matsuura K, Lefebvre PA, Kamiya R, Hirono M. (2004). Bld10p, a novel protein essential for basal body assembly in Chlamydomonas: localization to the cartwheel, the first ninefold symmetrical structure appearing during assembly. J Cell Biol. 165, 663–71.

Miller MS, Esparza JM, Lippa AM, Lux FG, Cole DG, Dutcher SK. (2005). Mutant kinesin-2 motor subunits increase chromosome loss. Mol. Biol. Cell 16, 3810–20.

Minami SA, Collis PS, Young EE, Weeks DP. (1981). Tubulin induction in C. reinhardtii: requirement for tubulin mRNA synthesis. Cell 24, 89–95.

Mohapatra L, Lagny TJ, Harbage D, Jelenkovic PR, Kondev J. (2017). The Limiting-Pool mechanism fails to control the size of multiple organelles. Cell Systems. 4, 559–567

Mueller J, Perrone CA, Bower R, Cole DG, Porter ME. (2005). The FLA3 KAP subunit is required for localization of kinesin-2 to the site of flagellar assembly and processive anterograde intraflagellar transport. Mol Biol Cell 16, 1341–54.

Mykles DL (2021). Signaling pathways that regulate the crustacean molting gland. Font. Endocrinol. 2: 674711

Pan J, Wang Q, Snell WJ. (2004). An aurora kinase is essential for flagellar disassembly in Chlamydomonas. Dev. Cell 6, 445–451.

Pan J, Snell WJ. (2005). Chlamydomonas shortens its flagella by activating axonemal disassembly, stimulating IFT particle trafficking, and blocking anterograde cargo loading. Dev. Cell 9, 431–8.

Periz G, Dharia D, Miller SH, Keller LR. (2007). Flagellar elongation and gene expression in Chlamydomonas reinhardtii. Eukaryot. Cell 6, 1411–20.

Pigino G, Geimer, S, Lanzavecchia, S., Paccagnini, E., Cantele, F., Diener, D.R., Rosenbaum, J.L., and Lupetti, P. (2009). Electron-tomographic analysis of intraflagellar transport particle trains in situ. J. Cell Biol. 187, 135–48.

Quarmby LM. (2004). Cellular deflagellation. Int. Rev. Cytol. 233, 47–91.

Qin, H., Diener, D.R., Geimer, S., Cole, D.G., and Rosenbaum, J.L. (2004). Intraflagellar transport (IFT) cargo: IFT transports flagellar precursors to the tip and turnover products to the cell body. J. Cell Biol. 164, 255–66.

Rafelski SM, Marshall WF. (2008). Building the cell: design principles of cellular architecture. Nat. Rev. Mol. Cell Biol. 9, 593–602.

Randall J. (1969). The flagellar apparatus as a model organelle for the study of growth and morphopoiesis. Proc. Roy. Soc. B. 173, 31–62.

Rosenbaum JL, Moulder JE, Ringo DL. (1969). Flagellar elongation and shortening in *Chlamydomonas*. The use of cycloheximide and colchicine to study the synthesis and assembly of flagellar proteins. J. Cell Biol. 41, 600–19.

Rosenbaum JL and Witman GB. (2002). Intraflagellar transport. Nat. Rev. Mol. Cell Biol. 3, 813–25.

Schloss JA, Silflow CD, Rosenbaum JL. (1984). mRNA abundance changes during flagellar regeneration in Chlamydomonas reinhardtii. Mol. Cell Biol. 4, 424–34.

Song L, Dentler WL. (2001). Flagellar protein dynamics in *Chlamydomonas*. J. Biol. Chem. 276, 29754–63.

Stolc V, Samanta MP, Tongprasit W, Marshall WF. (2005). Genome-wide transcriptional analysis of flagellar regeneration in Chlamydomonas reinhardtii identifies orthologs of ciliary disease genes. Proc. Natl. Acad. Sci. U.S.A. 102, 3703–7.

Walther Z, Vashishtha M, Hall JL. (1994). The Chlamydomonas FLA10 gene encodes a novel kinesin-homologous protein. J Cell Biol. 126, 175–88

Weeks DP, Collis P, Gealt MA. (1977). Control of induction of tubulin synthesis in Chlamydomonas reinhardtii. Nature 268, 667–8.

Wemmer KA, Marshall WF. (2007). Flagellar length control in Chlamydomonas - a paradigm for organelle size regulation. Int. Rev. Cytol. 260, 175–212.

Wemmer K, Ludington W, Marshall WF. (2020). Testing the role of intraflagellar transport in flagellar length control using length-altering mutants of Chlamydomonas. Philos Trans R Soc Lond B Biol Sci 375, 20190159

Wren KN, Craft JM, Tritschler D, Schauer A, Patel DK, Smith EF, Porter ME, Kner P, Lechtreck KF. (2013). A differential cargo-loading model of ciliary length regulation by IFT. Curr Biol 23, 2463–2471.

Wright RL, Chojnacki B, Jarvik JW. (1983). Abnormal basal-body number, location, and orientation in a striated fiber-defective mutant of *Chlamydomonas reinhardtii*. J. Cell Biol. 96, 1697–1707.

Zones JM, Blaby IK, Merchant SS, Umen JG. (2015). High-Resolution Profiling of a Synchronized Diurnal Transcriptome from Chlamydomonas reinhardtii Reveals Continuous Cell and Metabolic Differentiation. Plant Cell. 27, 2743–69.

